# Spatial Patterning of Laminin and N-Cadherin for Human Induced Pluripotent Stem Cell-Derived Cardiomyocytes (hiPSC-CMs)

**DOI:** 10.1101/2023.09.29.560252

**Authors:** Kerry V. Lane, Liam P. Dow, Erica A. Castilloa, Rémi Boros, Sam D. Feinstein, Gaspard Pardon, Beth L. Pruitt

**Author notes:** Equal contributions (Kerry V. Lane, Liam P. Dow).

## Abstract

Controlling cellular shape with protein micropatterning can mimic physiological morphologies and has been shown to improve reproducibility, enhancing our ability to collect statistics on single-cell behaviors. It has also advanced efforts in developing human induced pluripotent stem cell-derived cardiomyocytes (hiPSC-CMs) as a promising human model for studies of heart structure and function. hiPSC-CMs have key physiological differences from primary human cardiomyocytes (CMs), including lower sarcomere alignment and contractility, smaller area and lower aspect ratio, and lower force production. Protein micropatterning has been demonstrated to make hiPSC-CMs behave more like primary human CMs across these metrics. However, these micropatterned models typically use only extracellular matrix (ECM) proteins and have not investigated whether providing a protein associated with CM-CM interactions, such as N-cadherin, further enhances hiPSC-CM structure and function. Here, we developed a novel dual-protein patterning process to geometrically control single-cell CM placement on deformable hydrogels suitable for traction force microscopy (TFM). The patterns were comprised of rectangular laminin islands for attachment across the majority of the cell area, with N-cadherin “end-caps” imitating cell-cell interactions. We first photopatterned two proteins on a glass coverslip using a two-step process with photomolecular adsorption of proteins. After both photopatterning steps were complete, we transferred the pattern from the coverslip to a physiologically relevant ∼10-kPa polyacrylamide hydrogel. We seeded α-actinin-tagged hiPSC-CMs on the dual-protein-patterned hydrogels and verified interaction between the hiPSC-CMs and the N-cadherin end-caps via immunofluorescent staining. We found hiPSC-CMs on dual-protein patterns have a higher cell area and contractility in the direction of sarcomere organization than those on laminin-only patterns, but no difference in sarcomere organization or force production. While N-cadherin modestly improves the single-cell patterned hiPSC-CM model, it is not sufficient to replicate the role of cell-cell contacts in CM development for in vitro hiPSC-CM systems.

## Introduction

Human induced pluripotent stem cell derived cardiomyocytes (hiPSC-CMs) are a promising model to bridge the gap between human heart function and the studies of the human heart in animal models [1–4]. Developments in cardiomyocyte (CM) differentiation protocols have expanded the use of hiPSC-CMs in research [5] and engineering interventions have helped overcome limitations to the use of hiPSC-CMs as models for primary human CMs. These limitations include differences in structure and function between hiPSC-CMs and adult human CMs, such as CM morphology, sarcomere organization, and contractile force [4, 6].

Cardiac organoids or engineered heart tissues (EHTs) offer one approach to improving hiPSC-CM structure and function on average [7–10]. EHTs are commonly composed of hiPSC-CMs supported via some scaffolding, often made of extracellular matrix (ECM) proteins with mechanical properties similar to the myocardium [7]. EHTs mimic the 3D, multicellular environment of human CMs, and exhibit improved structure and function in hiPSC-CM – with sarcomere length and alignment similar to adult human CMs [9, 10]. However, EHTs require large numbers of hiPSC-CMs and supporting cells, show considerable heterogeneity in cell size and shape, and their complexity impedes live-cell imaging of subcellular structures, preventing investigations of sarcomere dynamics [6].

While EHTs provide a more realistic tissue environment for hiPSC-CMs, single hiPSC-CMs are prominently used in studies that seek to investigate intracellular processes, including assessments of cardiotoxicity in pre-clinical drug studies [11–14]. These studies use single-cell hiPSC-CMs to assess changes in contractile dynamics and ion channel function to investigate common and dangerous side effects of drugs before entering into clinical trials [11–14]. For this reason, developing better single-cell hiPSC-CMs is an important goal that could facilitate lower cost and higher throughput drug screening assays.

Protein micropatterning on hydrogels mimicking the mechanical properties of the myocardium provides an engineering approach to improve the structure and function of hiPSC-CM for single-cell assays [15, 16]. Protein micropatterning allows for the control of hiPSC-CM morphology by culturing the cells on rectangular ECM protein patterns in aspect ratios greater than or equal to 5:1, similar to the aspect ratio of adult human CMs [1, 15-17]. Patterned hiPSC-CMs present more highly aligned myofibrils and greater contractile forces than unpatterned hiPSC-CMs [16]. With these single-cell hiPSC-CMs, we have a high degree of control over the microenvironment and can study sarcomere dynamics and, by including fiducials in the deformable substrate, we can dynamically monitor force production.

Protein micropatterning has been used with a range of cell types to control cell shape and to create organized arrays of cells that allow for high-throughput imaging and analysis [18, 19]. Additionally, patterned cells have been shown to be more highly reproducible, creating more consistent intracellular phenotypes [18–20]. Théry and colleagues found that protein micropatterning of human retinal pigment cells created less intercellular variability and greater control over internal organization of the cells [18]. Tseng, et al. found that cell-cell junction positioning of mammary epithelial cells could be controlled by varying the spatial organization of ECM protein micropatterns [19]. Additionally, Rothenberg, et al. found that varying geometries of ECM protein micropatterns affected the number and organization of focal adhesions [20]. The reproducibility and higher control over subcellular organization are important benefits of the single-cell hiPSC-CM model, especially when screening for changing phenotypes, such as in drug studies [11, 12, 14].

Previous studies have investigated the effects of protein micropatterning on hiPSC-CMs, primarily focusing on the ECM proteins. A few studies have investigated CM-CM interactions using N-cadherin-functionalized substrates, culturing rat CMs on N-cadherin-coated polyacrylamide (PA) hydrogels [21] and N-cadherin-patterned glass [22]. Chopra, et al. coated PA hydrogels with N-cadherin, using anti-Fc antibody as a linking protein and stabilizing the proteins on the hydrogel with the crosslinker sulfo-N-sulfosuccini-midyl-6-(4’-azido-2’-nitrophenylamino) hexanoate (Sulfo-SANPAH) [21]. They then cultured single-cell rat CMs on the N-cadherin-coated hydrogels and found that the CMs produced traction force with a similar magnitude as single-cell rat CMs on ECM proteins [21]. Additionally, they showed that N-cadherin-mediated mechanotransduction is distinct from ECM-mediated mechanotransduction [21]. In another study, Chopra and colleagues patterned single-cell rat CMs on N-cadherin patterns on glass to learn that alpha-catenin serves as an adapter protein for N-cadherin-mediated mechanotransduction [22]. While these studies have provided useful insight into the role of N-cadherin in CM mechanotransduction, their translatability is limited because they are in rat CMs, which have key physiological differences compared to human CMs [23, 24].

Some studies have investigated the role of N-cadherin in the heart using single-cell, multicellular, and whole animal models from rats, mice, cats, and chicks [25–27]. Goncharova, et al. found that blocking N-cadherin inhibits the development and organization of sarcomeres in *in vitro* single-cell rat and chick CMs cultured on glass [25]. Additionally, Simpson and colleagues showed that cell-cell contacts are necessary for *in vitro* feline CMs cultured on laminin-coated petri dishes to stabilize myofibrils and to beat on their own [26]. In a whole animal study, Wu and colleagues showed that N-cadherin and integrin-mediated stabilization of myofibrils occur independently during development [27].

These previous *in vivo* and *in vitro* studies suggest that N-cadherin is relevant to CM structure, function, and development [21, 22, 25-27]. These studies used cells from animal models, limiting the relevance to the human heart. Additionally, all but one [21] used non-physiologic stiffness substrates and could not monitor CM contractile dynamics. Here, we bridge these gaps using single-cell hiPSC-CMs on protein-patterned deformable hydrogels suitable for traction force microscopy (TFM). The single-cell hiPSC-CM model allows us to investigate the impact of N-cadherin adhesions in human CM structure, including subcellular structures like sarcomeres, but also their contractile function.

Here, we asked whether pattering both laminin (an ECM protein secreted by CMs and abundant in their native microenvironment) and N-cadherin improves the structure and function of single-cell hiPSC-CMs. To answer this, we developed a method for consistent and precise dual-protein patterning on deformable hydrogels, and tested the effects of these substrates on single-cell hiPSC-CMs. N-cadherin is a relatively short (130 kDa), asymmetric protein which requires rotational freedom for proper conformation and binding [28]. To ensure the covalent attachment of functional N-cadherin, we adapted a protocol from Sarker, et al. [29] for the covalent attachment of Protein A as a linker to bind N-cadherin with an Fc-domain [30]. We used our protein micropatterning method to imitate both CM-ECM and CM-CM interactions for single-cell hiPSC-CMs. We hypothesized that utilizing the dual-protein patterning to imitate CM-ECM and CM-CM interactions would improve hiPSC-CM structure, *e.g.,* cell spread area, sarcomere alignment, and contractile function, *i.e.*, sarcomere contractility and force production.

## Results and Discussion

We created two single cell patterns: i) single protein patterns consisting of laminin rectangles, and ii) dual protein patterns, consisting of laminin rectangles flanked by N-cadherin caps (Figure 1a). We patterned arrays of alternating single and dual protein patterns on each coverslip. We then transferred the protein patterns to a ∼6.8-kPa polyacrylamide (PA) hydrogel. We verified the patterning and the pattern transfer to PA hydrogel using a laminin antibody and a pan-cadherin antibody (Figure 1a). Finally, we seeded hiPSC-CMs on the patterns (Figure 1b) and assessed their morphology, force production, and sarcomere organization and contractility on the laminin-only and dual-protein patterns.

**Figure 1.**
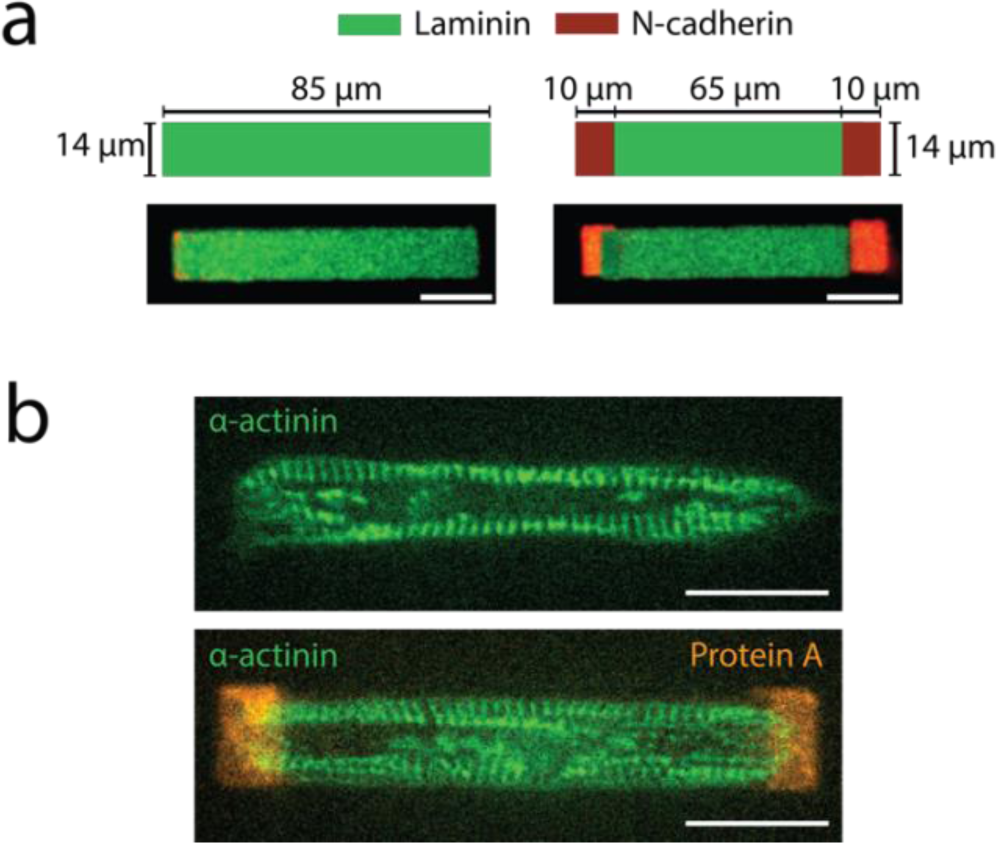
Dual-protein and laminin-only patterns. **(a)** Schematics and representative images of fluorescently labelled laminin-only and dual-protein patterns on PA hydrogels. Green is laminin and red is N-cadherin. **(b)** Representative image of hiPSC-CMs on laminin-only (top) and dual-protein (bottom) patterns, with sarcomeres (green) and Protein A (orange) visible. Scale bars are 20 μm.

### Mechanical characterization of polyacrylamide hydrogels

We used atomic force microscopy (AFM) to characterize four polyacrylamide hydrogels (Figure 2). Using the Hertz model (Figure 2a), we found the mean stiffness of the polyacrylamide hydrogels to be 6.8 kPa, with a standard deviation of 1.5 kPa.

**Figure 2.**
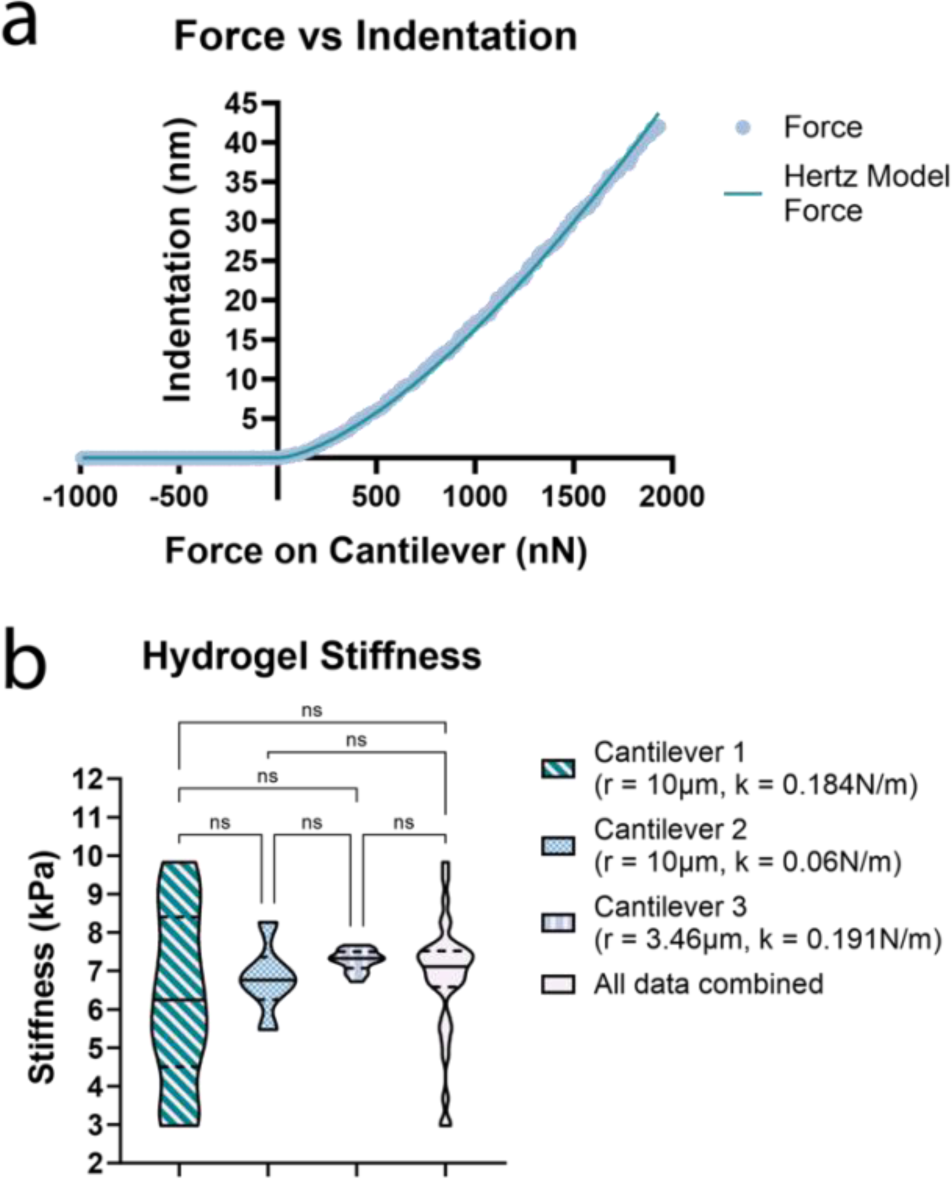
Mechanical characterization of polyacrylamide hydrogels. **(a)** Representative example of a force-indentation curve for a ∼6.8 kPa hydrogel, with measured force (light blue circular markers) plotted alongside the Hertz model calculated force (teal line). **(b)** PA hydrogel stiffness measurements with each cantilever. For stiffness data, n = 31. In (b), centerlines indicate medians, dotted lines indicate 25^th^ and 75^th^ percentiles.

We used multiple AFM cantilevers due to PA hydrogel build-up on cantilever tips after ∼10-20 measurements. The three cantilevers had the following properties: (1) r_t_ = 10μm, k_c_ = 0.184N/m, sensitivity = 0.047V/nm, (2) r_t_ = 10μm, k_c_ = 0.060N/m, sensitivity = 0.027V/nm, and (3) r_t_ = 3.46μm, k_c_ = 0.191N/m, sensitivity = 0.042V/nm.

We compared the data from each cantilever using a Kruskal–Wallis one-way analysis of variance test to ensure that the results were consistent. There was no statistically significant difference between any of the data sets, including the data set containing all of the datapoints together (p-values between 0.44 and >0.9999; Figure 2b).

### Validation of hiPSC-CM interaction with N-cadherin on dual-protein patterns

We first sought to verify that the hiPSC-CMs were interacting with the N-cadherin end-caps on the dual-protein patterns. We stained hiPSC-CMs on dual-protein patterns with a pan-cadherin primary antibody and AlexaFluor-647 secondary antibody. We manually outlined each cell in FIJI (ImageJ) as described in *Methods*. We outlined the N-cadherin end-cap using the 647 channel (N-cadherin). We isolated the overlap between the cell outline and the N-cadherin end-cap and quantified the average fluorescence intensities in the overlap area and the N-cadherin end-cap area (Figure 3a).

**Figure 3.**
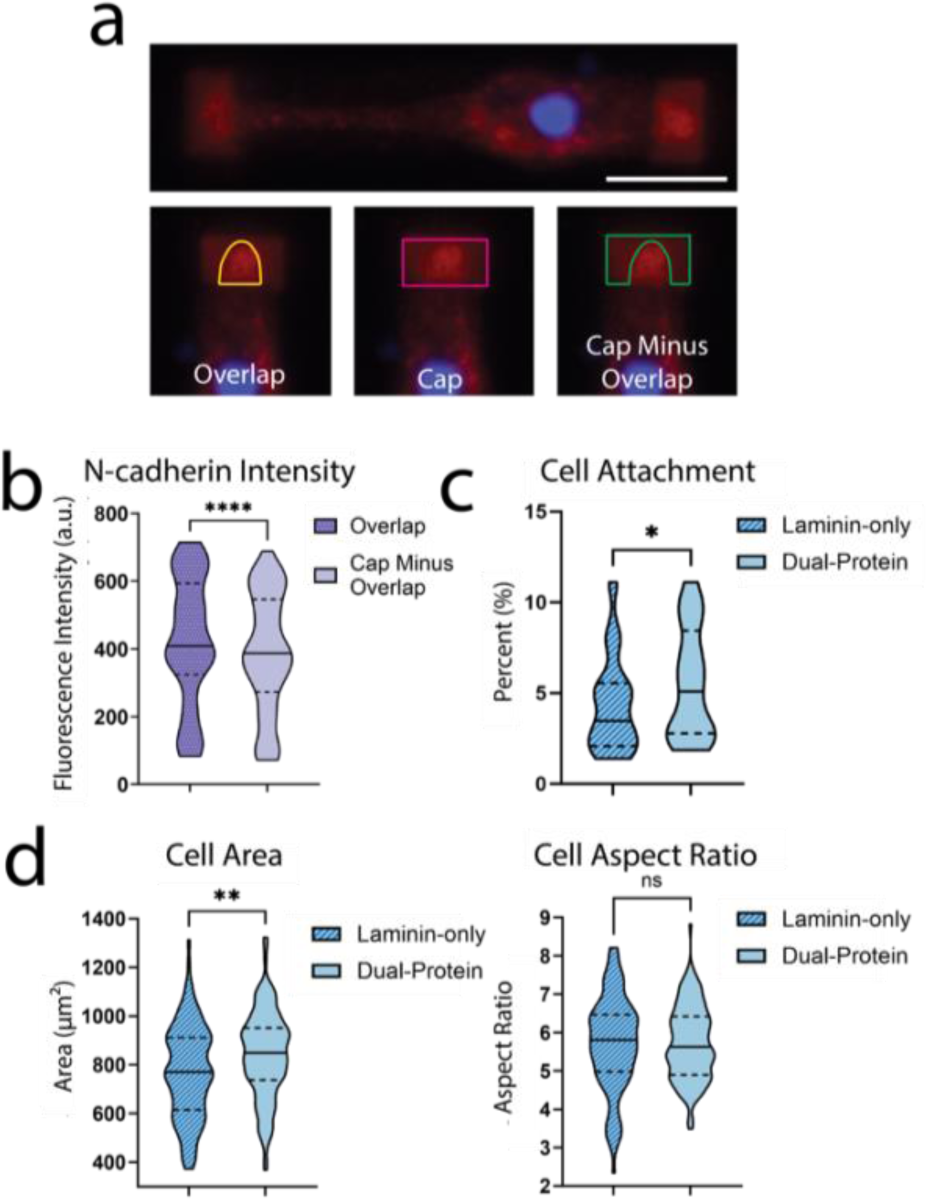
Dual-protein-patterned hiPSC-CMs have greater cell area than those on laminin-only patterns. **(a)** Verification of hiPSC-CM interaction with N-cadherin. hiPSC-CM stained for N-cadherin (red) and cell nucleus (blue). Overlap of cell and N-cadherin measured using the outlines shown and **(b)** the resulting fluorescent intensities of the overlapping area and the end-cap area without the overlap are visualized in the plot to the right. For N-cadherin verification data, n = 62. **(c)** hiPSC-CM cell area and aspect ratio and **(d)** attachment rates of hiPSC-CMs on laminin-only (right) and dual-protein (left) patterns. For cell area, aspect ratio, and attachment data, n = 116 for laminin-only patterned condition and n = 131 for dual-protein patterned condition. For all plots, centerlines indicate medians, dotted lines indicate 25^th^ and 75^th^ percentiles. P-values with significance at P<0.05 are designated with (*), P<0.005 are designated with (**), P<0.0005 are designated with (***), and P<0.0001 are designated with (****). Scale bar is 20 μm.

We compared the two conditions using a two-tailed Wilcoxon matched-pairs signed rank test. We observed a significant difference (p < 0.0001) between the intensity of N-cadherin signal where the hiPSC-CMs overlapped the N-cadherin patterns and the intensity of the N-cadherin patterns themselves (Figure 3b). This overlap confirms that the hiPSC-CMs localized endogenous N-cadherin on the *in vitro* N-cadherin patterns.

### hiPSC-CMs on dual-protein patterns have increased cell area and attachment rates

To investigate the effect of dual-protein patterns on hiPSC-CM structure, we assessed cell area and aspect ratio, as well as the rate of attachment of hiPSC-CMs to laminin-only and dual-protein patterns.

Most studies that pattern hiPSC-CMs use rectangles with an aspect ratio of 5:1 to 7:1, corresponding to the range of adult human ventricular CMs [1, 16, 17]. In early experiments, we used patterns with an aspect ratio of 7:1, but found most cells did not fill the entire pattern, meaning many did not reach the N-cadherin end-caps on the dual-protein patterns. We adjusted the length of the patterns to address this issue, bringing our pattern aspect ratio down to ∼6:1.

The cell area and aspect ratio of the hiPSC-CMs were assessed by drawing an outline of the cell in FIJI, as described in *Methods*. The areas of hiPSC-CMs on dual-protein patterns were larger than those of laminin-only patterns, with average areas of 824.4μm^2^ and 739.9μm^2^, respectively (P = 0.0023; Figure 3d). For reference, the total pattern area is 1190μm^2^.

The mean aspect ratios of hiPSC-CMs on the laminin-only and dual-protein patterns were 5.4 and 5.5 respectively (Figure 3d). The laminin-only-patterned hiPSC-CMs had a larger variance than the dual-protein-patterned hiPSC-CMs, with standard deviations of 1.21 and 0.96, respectively (F-test p = 0.0085). The mean aspect ratios were compared using an unpaired, two-tailed T-test with Welch’s correction and had no significant difference (P = 0.8705). These results confirm that both laminin-only and dual-protein patterns support spreading of hiPSC-CM near the patterned aspect ratio. In addition to hiPSC-CM area and aspect ratio, we considered whether the addition of N-cadherin end-caps influenced the rate of cell attachment. To determine attachment rates, we counted the number of single-cell hiPSC-CMs attached to each pattern type. For this data, we included all cells on dual-protein patterns, regardless of whether they overlapped an N-cadherin end-cap or not. We assessed the difference in attachment rates using a parametric, two-tailed, ratio paired T-test. The rate of attachment was higher on dual-protein patterns than laminin-only patterns, with average attachment rates of 5.5% and 4.3%, respectively, on seventeen total devices (P = 0.0441; Figure 3c). This increased attachment suggests that dual-protein patterns may promote increased hiPSC-CM attachment rates, even if the hiPSC-CMs didn’t overlap the N-cadherin end-caps at the time of fixation/imaging.

### hiPSC-CMs on dual-protein patterns have increased contractility in the direction of sarcomere alignment

To investigate the effect of dual-protein patterns on hiPSC-CM function, we analyzed the contractility of the hiPSC-CM sarcomeres. Sarcomeric contractility is a commonly used metric to compare hiPSC-CMs to adult human CMs, with adult human CMs exhibiting greater contractility than hiPSC-CMs [1, 4, 16].

We analyzed sarcomere contractility and organization using a previously published, open-source program called Sarc-Graph [31]. Sarc-Graph segments videos of fluorescently-tagged sarcomeres in beating hiPSC-CMs and outputs parameters representing the orientation, spacing, and contractility of the sarcomeres, all of which are important metrics for hiPSC-CM structure and function [31].

In overall sarcomere shortening, we did not see a significant difference between hiPSC-CMs on laminin-only vs dual-protein patterns (Figure 4a). The percent shortening is calculated by taking the difference between the maximum and minimum length for an individual sarcomere and dividing it by the average length of the sarcomere [31]. The percent sarcomere shortening for all of the sarcomeres in one cell are averaged to calculate a single percent sarcomere shortening for each hiPSC-CM. The mean percent sarcomere shortening for laminin-only and dual-protein patterns were not significantly different, with values of 16.89% and 17.67%, respectively (P = 0.6273). To further investigate sarcomere contractility, we assessed *C_||_*, a parameter calculated by Sarc-Graph that relates sarcomere contractility to sarcomere alignment and represents the shortening of the entire cell domain in the direction of dominant sarcomere orientation [31].

**Figure 4.**
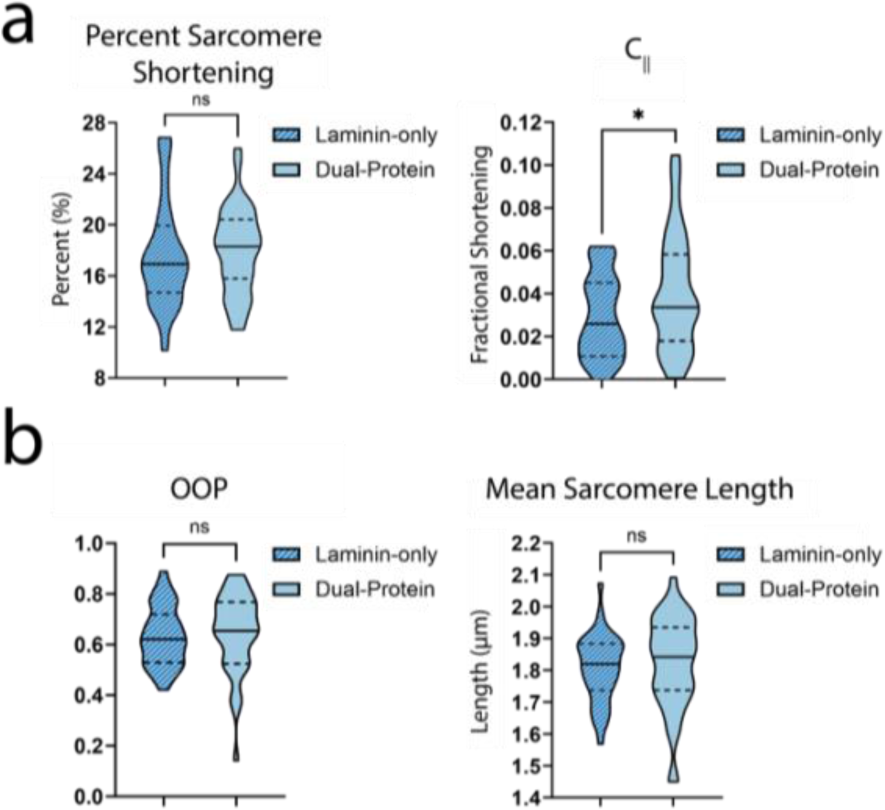
Dual-protein-patterned hiPSC-CMs have higher contractility in the direction of sarcomere organization, but no difference in sarcomere alignment compared to laminin-only-patterned hiPSC-CMs. **(a)** Overall percent sarcomere shortening and fractional sarcomere shortening in the direction of sarcomere organization (C_||_) for hiPSC-CMs on laminin-only and dual-protein patterns. **(b)** OOP and mean sarcomere length for hiPSC-CMs on laminin-only (right) and dual-protein (left) patterns. For both laminin-only and dual-protein patterned conditions, n = 39. For all plots, centerlines indicate medians, dotted lines indicate 25^th^ and 75^th^ percentiles. P-values with significance at P<0.05 are designated with (*).

To define *C_||_*, we first need to understand how Sarc-Graph defines sarcomere alignment. Zhao, et al. use a structural tensor to quantitatively assess the sarcomere orientation, defining the structural tensor with the equation

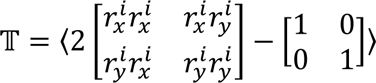

where **r** = [*r^i^_x_*, *r^i^_y_*] is a unit vector representing the orientation of the i^th^ sarcomere [31]. The structural tensor has eigenvalues of *a*_max_ and *a*_min_, with *a*_max_ providing the magnitude of sarcomere alignment (OOP). The eigenvectors of the structural tensor are **v**_max_ and **v**_min_, with **v**_max_ representing the direction of sarcomere alignment [31].

To define the contractility, Zhao and colleagues define the deformation gradient **D**_avg_ with the equation

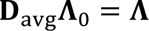

where **Λ**_0_ = [**v**_01_, **v**_02_,…, **v**_0*n*_], representing the vectors **v** that connect neighboring sarcomeres in the initial reference frame, and **Λ** = [**v**_1_, **v**_2_,…, **v***_n_*], representing the vectors **v** connecting neighboring sarcomeres in the current deformed frame [31].

Zhao, et al. relate the deformation gradient back to the sarcomere alignment expressions with the equation **v**_max_ = **D**_avg_**v**_0_ [31], where **v**_max_ is the eigenvector representing the direction of sarcomere alignment in the contracted (deformed) state and **v**_0_ representing the direction of sarcomere alignment in the relaxed (initial) state [31]. They then define *C_||_* as

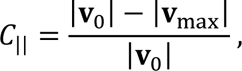

representing the fractional shortening of the sarcomeres in the direction of sarcomere alignment [31].

We found that hiPSC-CMs on dual-protein patterns had larger *C*_||_ values, with an average of 0.041 compared to 0.029 for laminin-only-patterned hiPSC-CMs (P = 0.0294; Figure 4a). This result suggests that the contractility of the sarcomeres along the long axis of the myofibrils is enhanced in hiPSC-CMs patterned with N-cadherin end-caps mimicking CM-CM interactions.

We determined the statistical significance with an unpaired T-test with a Welch’s correction for the unequal variance (F-test P = 0.0403) in the dual-protein-patterned data (standard deviation = 0.026) compared to the laminin-only-patterned data (standard deviation = 0.019).

### hiPSC-CM sarcomere organization is similar on laminin-only and dual-protein patterns

We further examined the impact of dual-protein patterning on hiPSC-CM structure by analyzing sarcomere structure and organization. Previous studies have suggested that adult CMs have average sarcomere lengths of ∼2.2 μm, while immature CMs have average sarcomere lengths of ∼1.6 μm [1].

To investigate sarcomere structure and organization, we assessed hiPSC-CM sarcomere lengths and the orientational order parameter (OOP), a metric commonly used to assess sarcomere organization in CMs that ranges from zero (random orientation) to 1 (perfectly aligned) [31]. We hypothesized that hiPSC-CMs on dual-protein patterns would have more highly organized sarcomeres because previous studies found that N-cadherin is important in sarcomere formation and organization [21, 22, 25-27, 32]. Surprisingly, sarcomeric organization, as assessed via the OOP, was not significantly different between hiPSC-CMs on laminin-only and dual-protein patterns, with average values of 0.6330 and 0.6254, respectively (P = 0.6205; Figure 4b). The lack of difference in sarcomere organization between laminin-only and dual-protein patterned hiPSC-CMs suggests that the patterned N-cadherin end-caps are not sufficient to increase organization.

We also found no difference between the mean sarcomere lengths of hiPSC-CMs on laminin-only and dual-protein patterns. Both groups had mean sarcomere lengths of 1.8 μm, with minimum and maximum sarcomere lengths of 1.7 μm and 2 μm, respectively (P = 0.6193; Figure 4b). We assessed the statistical significance using an unpaired T-test with a Welch’s correction because the standard deviations of the laminin-only and dual-protein data, 0.10 μm and 0.14 μm, respectively, were significantly different (F-test p = 0.0408).

### hiPSC-CM force production is similar on laminin-only and dual-protein patterns

To further investigate the impact of dual-protein patterning on hiPSC-CM function, we assessed force production of hiPSC-CMs with Traction Force Microscopy (TFM), using the streamlined TFM module of a custom, open-source code called CONTRAX [33, 34]. The streamlined TFM module of CONTRAX provides a user-friendly TFM analysis tool that reads in fluorescent microbead displacement videos and assesses a number of functional metrics, including traction force [33–35].

We looked at multiple parameters output by CONTRAX, including total force production, peak traction stress, average contraction displacement, and contraction velocity. We also assessed total contractile moment, which is a scalar value representing the sum of moments taken at the center of the cell [36], and total impulse, which is the integrated area under the curve from the force versus time plot [33].

Force production, average contraction displacement, and contraction velocity are all directly related to hiPSC-CM contractility and are commonly used to assess CM function [16, 37, 38]. The peak traction stress avoids the homogenization caused by integrating over the cell area, providing insight into the maximum stress produced by each cell.

Total force production is calculated by integrating the traction stresses over the cell area at each time point, creating a trace of force versus time, and identifying the maximum amplitude of the trace peaks [33–35]. To assess average contraction displacement, CONTRAX averages the displacement magnitudes over the cell area for each video frame. CONTRAX then creates a one-dimensional trace of average displacement over time and extracts the displacement between the fully contracted and fully relaxed states [33, 34]. CONTRAX calculates the contraction velocity using the same displacement versus time trace, dividing the contraction displacement by the time elapsed during contraction [33, 34]. Peak traction stress is extracted by identifying the timepoint with greatest traction stress and reporting the highest absolute value of traction stress within that timepoint.

The total contractile moment provides information about the distribution of stresses by weighting the traction forces by their distance from the cell center. We hypothesized that the force production of hiPSC-CMs might be more concentrated at the N-cadherin end-caps compared to the hiPSC-CMs on laminin-only patterns. The contractile moments are calculated by multiplying the traction force by the distance from the cell center, using the equation

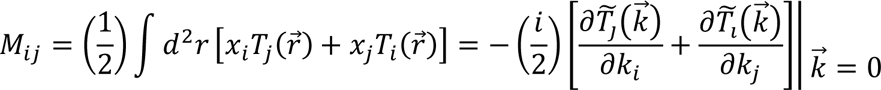

where 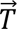(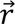) is the traction vector at point 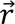 on the hydrogel surface [36]. The second expression represents the equation in Fourier space, which we operate in when using Fourier Transform Traction Cytometry (FTTC; see *Methods*). The tilde indicates the two dimensional Fourier transform with wave vector 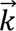[36]. Using *x* and *y* for indices *i* and *j*, we have four possible equations representing scalar values: *M_xx_*, *M_yy_*, *M_xy_*, and *M_yx_*. *M_xy_* and *M_yx_* represent torque on the substrate due to the traction forces, and *M_xx_* and *M_yy_* represent contractile forces weighted by their distance to the center of the cell [36]. To investigate the weighted contribution of *x*- and *y*-forces to the cell contraction, we assess the total contractile moment by adding the *x*- and *y*-contractile moments together: *μ* = *M*_*xx*_ + *M*_*yy*_ [36].

The total impulse is calculated by calculating the area under the force versus time trace [33, 39]. It is equivalent to the tension-integral parameter used in a previous study that found that hypertrophic and dilated cardiomyopathies could be differentiated in hiPSC-CMs by the tension-integral magnitude [39]. We did not expect to see a difference in total impulse between our two conditions, but assessed the parameter to verify that there was no difference.

For all of these parameters except for average contraction displacement and contraction velocity, we used the peak measurements, meaning the values calculated when the cell is either fully contracted or fully relaxed. Our analysis tools assume these are quasi-static states; however, we note this assumption is violated during active contraction or relaxation occurring across all frames of our ∼800 frame videos. We did not find significant differences between total force, peak stress, average contraction displacement, contraction velocity, total contractile moment, or total impulse (Figure 5). This result is consistent with the overall contractility result, further indicating that the difference in functional outputs of hiPSC-CMs on laminin-only and dual-protein patterns are modest.

**Figure 5.**
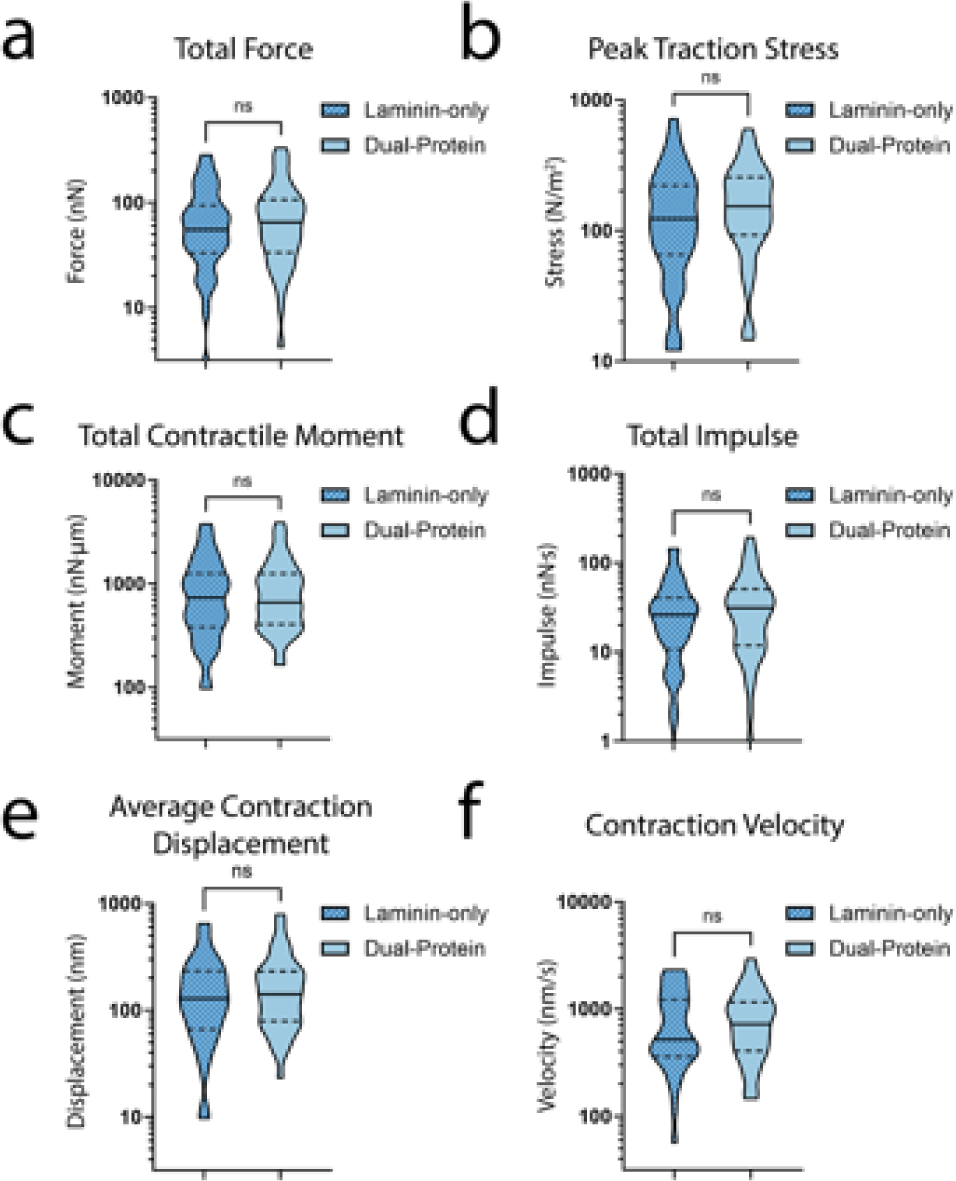
Dual-protein-patterned hiPSC-CMs demonstrate no difference in **(a)** total force, **(b)** peak traction stress, **(c)** total contractile moment, **(d)** total impulse, **(e)** average contraction displacement, and **(f)** contraction velocity compared to laminin-only-patterned hiPSC-CMs. For all data in this figure, N = 81 with n = 35 for laminin-only and n = 46 for dual-protein condition. For all plots, centerlines indicate medians, dotted lines indicate 25^th^ and 75^th^ percentiles.

Within both laminin-only and dual-protein pattern conditions, we saw a large variance in force production, as can be seen in Figure 5a. For the laminin-only-patterned hiPSC-CMs, the mean force was 104.1nN +/-98.2nN, while the dual-protein-patterned hiPSC-CMs had an average force of 118.3nN +/-107.1nN.

All of the parameters reported by CONTRAX, including those assessed above, are single scalar values for each cell. These parameters allow for quantification of force production but result in the loss of spatial information of hiPSC-CM contractility. This flattening of spatially distributed forces into scalar values is necessary to quantify and assess data sets that consist of ∼800 frames per video and hundreds of displacement data points in each frame of each video. While the scalar outputs provide quantifiable comparisons between cells, the spatial information can contain differences in force distribution that are not discernible in scalar variables. To investigate the spatial distribution of force production for each condition, we averaged the peak contraction traction stress heatmaps of all the cells in each condition (Figure 6), with 34 cells in the laminin-only condition and 46 cells in the dual-protein condition. We visually see a difference between the traction stresses in the laminin-only- and dual-protein-patterned conditions, with the peak stresses appearing to occur closer together in dual-protein-patterned hiPSC-CMs compared to laminin-only-patterned hiPSC-CMs. The inward shift in peak traction stress location for the hiPSC-CMs on dual-protein patterns could be reflecting the inward shift of the laminin pattern boundary compared to the laminin-only patterns. The red squares in Figure 6a indicate the location of the N-cadherin end-caps and the peak traction stresses occur within the laminin portion of the dual-protein patterns. This suggests that the focal adhesions still drive the traction force production in hiPSC-CMs on dual-protein patterns. Additionally, we see lower traction stresses in the laminin-only-patterned hiPSC-CMs compared to the dual-protein-patterned hiPSC-CMs.

**Figure 6.**
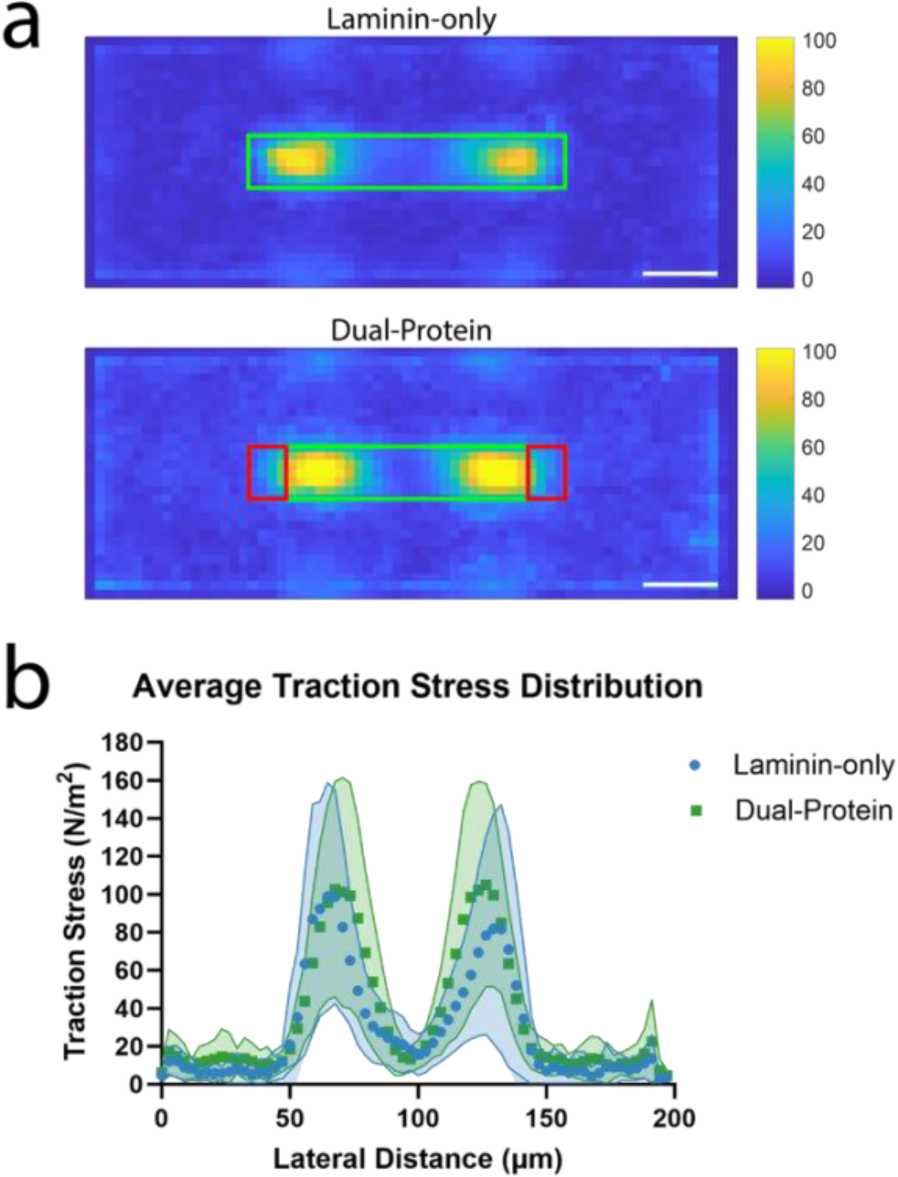
Spatial distribution of average traction stresses produced by hiPSC-CMs on laminin-only and dual-protein patterns. **(a)** Peak contraction traction stress heatmaps averaged over all cells in both laminin-only (n = 34) and dual-protein (n = 46) conditions. Green rectangles represent an estimation of the locations of laminin patterns, red rectangles represent an estimation of the locations of N-cadherin patterns. Color bars are in units of Pascals. **(b)** Mean traction stress distribution with respect to lateral distance, averaged over a ∼10 μm width in the center of each heatmap. The shaded areas represent the standard deviations. Scale bars are 20 μm.

While the location of peak traction stress appears to have moved toward the cell center, the area of the peak traction stress is larger in dual-protein-patterned hiPSC-CMs compared to laminin-only-patterned hiPSC-CMs. The areas of peak traction stress for the laminin-only patterns on the left and right are 170 μm^2^ and 155 μm^2^, respectively. For the dual-protein patterns, the areas of peak traction stress on the left and right are 310 μm^2^ and 292 μm^2^, respectively. This could suggest that while the mechanotransduction occurs near the edge of the laminin pattern, the hiPSC-CMs on dual-protein patterns are establishing mechanical connections to the patterns with larger areas.

## Conclusions

In this work, we have developed a method for precise patterning of multiple proteins on a single device and applied this method to create patterns mimicking cell-cell and cell-ECM interactions for hiPSC-CMs. Our method applies a novel dual patterning method to functional cellular studies. We have demonstrated the use of the method by making dual-protein patterns to mimic cell-cell interactions for single-cell hiPSC-CMs. Our results indicate that our dual-protein patterning increases hiPSC-CM attachment rates, spread area, and the efficiency of sarcomere contraction along myofibrils. We found no significant difference in force production, overall sarcomere contractility, and sarcomere organization between laminin-only and dual-protein patterned hiPSC-CMs, suggesting that N-cadherin end-caps offer modest benefit alone. More complexity is likely necessary to improve hiPSC-CM structure and function beyond single ECM patterning alone.

In addition to greater complexity in the patterning approach, more detailed spatial analysis of force production and sarcomere contractility could provide more insight into the differences between laminin-only and dual-protein-patterned hiPSC-CMs. Detailed spatial analysis is made difficult by the high quantity of data – there are almost 2,000 data points per frame, 800 frames per cell, and ∼40 cells per condition. In addition to the high volume of data, the data are heterogeneous, with variations in the time point of peak contraction for each cell as well as variations in cell area and cell location in relation to the pattern.

## Methods

### Protein patterning glass coverslips

N-cadherin is an asymmetric protein and a linking protein is necessary to ensure the extracellular binding site is available when patterning N-cadherin [30, 40, 41]. In this work, we used an N-cadherin Fc chimera (R&D Systems, 1388-NC-050) and a 546-nm fluorescently tagged Protein A (ThermoFisher, P11049) as our linking protein. Protein A is a protein with a high affinity to Fc-regions of IgG molecules [42]. It has been previously used to ensure the correct orientation of E-cadherin [41, 43]. The Fc region of the N-cadherin chimera protein binds to the Protein A, ensuring the correct orientation of the N-cadherin on the device surface.

To begin protein micropatterning, we activated 18mm diameter #1 glass coverslips with oxygen plasma for 5 minutes at 18W (Harrick, PDC-32G). Immediately after plasma treatment, we sealed an 8mm inner diameter silicone ring (B&J Rubber Products) to the center of the glass coverslip. The silicone ring was cut from a Silhouette CAMEO 3 electronic desktop cutter (Silhouette America). We immediately pipetted a solution of 100 µg/mL of poly(l-lysine)-graft-poly(ethylene glycol) (PLL(20)-g[3.5]-PEG(2); SuSoS AG) diluted in phosphate buffered saline (PBS; Gibco, ThermoFisher, 10010049) within the ring and incubated for 1 hour at room temperature. We rinsed the PLL-g-PEG thoroughly (10x) with PBS prior to micropatterning. Following PLL-g-PEG incubation and rinsing, we pipetted 20μL of UV sensitive photoinitiator (PLPP; Alvéole) into the silicone ring on the glass coverslip. Then we placed the glass coverslip on the stage of a Leico Dmi8 epifluorescence microscope equipped with a Fluotar 20x/0.40 NA objective and the Alvéole Primo photopatterning system (Alvéole) with a 375 nm, 7.10 mW laser. We made digital masks for the protein A “end-cap” patterns as well as the main laminin patterns using the open-source software Inkscape (https://inkscape.org). We used the pixel-to-micron ratio generated by Primo calibration to define the geometries of all patterns. We defined two patterns for constraining the single-cell hiPSC-CM: 1) 14 µm x 85 µm laminin-only patterns; and 2) 14 µm x 85 µm dual-protein patterns. Dual protein patterns are comprised of Protein A end-cap patterns (14 µm x 10 µm) overlapped with laminin rectangles (14 µm x 69 µm) by ∼10% to mitigate any alignment artifacts.

We loaded the digital masks into the Leonardo plugin (Alvéole Laboratory) on Micro-Manager software [44], and made a 6 by 6 array with 150 µm spacing between each instance of the patterns. We illuminated the glass coverslip with the first digital mask for the Protein A end-caps at a dosage of 1,000 mJ/mm^2^. Following micropatterning, we rinsed off the photo initiator with PBS and incubated the glass coverslip with 100 µL of a 100 µg/mL solution of 546-nm fluorescently tagged Protein A overnight at 4 °C. We then thoroughly rinsed the Protein A from the glass coverslip using PBS and added another 20 µL of PLPP photoinitiator. We illuminated the glass coverslip with the second digital mask for the laminin rectangles at a dosage of 1,000 mJ/mm^2^. We designed the array of patterns to alternate between dual-protein and laminin-only patterns. Before UV illumination, we aligned the shortened laminin bodies within the digital mask to the already-patterned protein A end-cap patterns using a Texas Red fluorescent excitation filter. Following this second UV illumination step, we rinsed off the photoinitiator with PBS and incubated the glass coverslip with a 500 µg/mL solution of laminin (Corning, 354232) for 2 hours at room temperature. For pattern verification experiments, we incubated the glass coverslip with a 500 µg/mL solution of green fluorescent laminin (Cytoskeleton, Inc., LMN02) for 2 hours at room temperature. Finally, we rinsed the patterns with PBS and removed the silicone containment ring prior to gel transfer. A schematic of the protein micropatterning process flow can be seen in Figure 7a.

**Figure 7.**
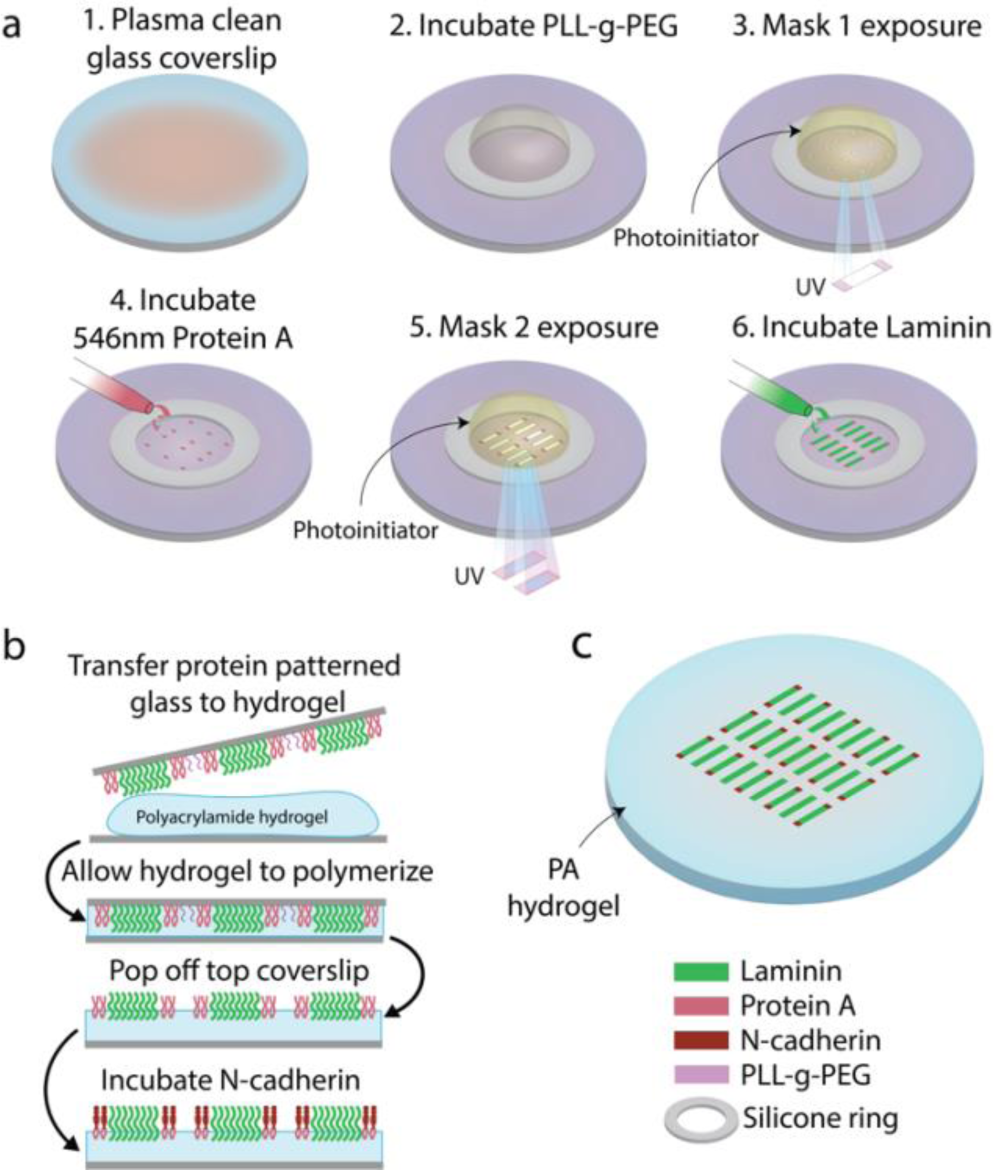
Dual-protein patterning of PA hydrogels. **(a)** Process flow of dual-protein patterning on a glass coverslip using photomolecular adsorption. **(b)** Process flow of transfer of dual-protein pattern from glass coverslip to PA hydrogel and incubation of N-cadherin. **(c)** Schematic of final result – alternating laminin-only and dual-protein patterns on PA hydrogel.

### Preparation of polyacrylamide (PA) hydrogels

In preliminary work, the Protein A—N-cadherin complex could not anchor hiPSC-CMs on unfunctionalized polyacrylamide (PA) hydrogels (Figure S2), so we employed oxidized N-Hydroxyethyl acrylamide (oHEA) to create a covalent bond between the hydrogel and the Protein A. We transferred the laminin and Protein A patterns to a ∼10kPa oHEA-functionalized PA hydrogel before adding N-cadherin, to ensure that the final hydrogel presented the N-cadherin binding domain on the hydrogel surface. We prepared the oHEA-functionalized PA hydrogels following a previously published protocol with some modifications [29]. Briefly, we began by oxidizing N-Hydroxyethyl acrylamide (HEA; Sigma, 697931) by adding 0.01 g of sodium metaperiodate (Sigma, 71859) to 2.338 mL of HEA, then incubating in the dark on a shaker for 4 hours. To adhere the PA hydrogels to the glass bottom dishes used in this work, we treated the glass with bind-silane. We prepared a solution with 95 μL of 100% ethanol, 50 μL of acetic acid, and 3μL of 3-(Trimethoxysilyl)propyl methacrylate (Bind-silane; Sigma, M6514). Then, we treated the glass with oxygen plasma for 30 seconds at 18W (Harrick, PDC-32G). Directly after plasma treating the glass, we added ∼50 μL of the bind-silane solution to the glass surface. We incubated the solution on the glass for 1 minute, after which we removed the excess bind-silane solution. We left the remaining solution to react for 10 minutes, after which we rinsed the glass twice with 1 mL of 100% ethanol and dried with nitrogen gas.

To prepare the PA hydrogel solution, we combined 732 μL of 40% Acrylamide solution (Bio-Rad, 1610140) and 260 μL of 2% Bis-acrylamide solution (Bio-Rad, 1610142) with 4.008 mL MilliQ water. For experiments used in TFM analysis, we added 326μL of 1.0 μm-diameter blue fluorescent microbeads (ThermoFisher, F8814) and decreased the MilliQ volume to 3.682 mL to maintain the total volume of 5mL. Finally, we added 200μL of oxidized HEA, bringing the total volume to 5.2 mL. Separately, we prepared a 10% weight by volume (w/v) solution of ammonium persulfate (APS; Sigma, A9164) in MilliQ water.

To begin polymerization, we added 2.6 μL of N,N,N′,N′-tetramethylethylenediamine (TEMED; Sigma, 411019) and 260 μL of the 10% w/v APS solution to the PA solution. We gently mixed the solution with a P1000 pipette and then pipetted 35μL of the solution onto the bind-silane-treated glass surface. We then placed a protein patterned coverslip on top of the solution, sandwiching the PA solution between the bind-silane-treated glass and the protein patterned glass. After casting, the hydrogel polymerized in the dark for 30 minutes before we hydrated it with PBS and left it to fully polymerize at 4°C for 6-8 hours. After full polymerization, we removed and discarded the protein patterned top coverslip. A schematic of the protein transfer to oHEA-functionalized PA hydrogel can be seen in Figure 7b.

Following the removal of the top coverslip, we aspirated the PBS from the dish and incubated each PA hydrogel with ∼50 μL of 100 μg/mL N-cadherin (R&D Systems, 1388-NC-050) for 3 hours at 4°C. After 3 hours, we washed the PA hydrogel three times with 1mL PBS and then stored it overnight at 4°C in PBS with 10% Antibiotic-Antimycotic 100X (Anti-Anti; Gibco, ThermoFisher, 15-240-062) and 1% bovine serum albumin (BSA; ThermoFisher, PI37525). The following day, we washed the hydrogel three times with 1 mL PBS and then stored it in PBS + 10% Anti-Anti until cell seeding. A schematic of the final device can be seen in Figure 7c.

We verified the patterning and the pattern transfer to PA hydrogel using green fluorescent laminin (as described in *Protein patterning glass coverslips*) and a pan-cadherin primary antibody (Sigma, C3678). We diluted the pan-cadherin antibody 1:200 in PBS and incubated on the devices for 1 hour at room temperature. We washed the devices three times with PBS and then incubated the devices with anti-rabbit AF-647 diluted 1:500 in PBS for 1 hour at room temperature. We rinsed the devices three times with PBS and then imaged (Figure 1a).

### Stiffness characterization by Atomic Force Microscopy (AFM)

We characterized our PA hydrogel stiffness using AFM indentation based on a previously published protocol [45]. Briefly, we used a WITec AFM (Alpha300) and large tip cantilevers with tip radii of curvature (r_t_) of 3.46 μm and 10 μm and nominal spring constants (k_c_) of 0.191 N/m (3.46 μm tip – Bruker, SAA-HPI) and 0.184 N/m and 0.191 N/M (10 μm tip – Bruker, SAA-SPH-10UM and MLCT-SPH-10UM). We measured four hydrogels, assessing 3 locations per hydrogel with between 2-3 measurements per location. Each hydrogel was measured 2 days after polymerization.

The PA hydrogels were attached to a glass bottom dish and submerged in PBS. We measured Optical Lever Sensitivities [46] before each experiment by performing force-distance scans against the glass surface of a glass bottom plate using the following parameters: feedback control with 1.0 V set point, 1% p-gain, and 0.2% i-gain, force-distance using 0.2 μm pull and 0.6 μm push at 0.2 μm/s speed.

After determining the Optical Lever Sensitivity, we loaded the PA hydrogel sample and centered the cantilever above a point on the hydrogel. We then approached the surface using the following parameters: 1.0 V set point, 1% p-gain, and 0.2% i-gain. We then performed the force-distance curve measurement using WITec’s Distance Curve mode with approach and retract distances of 20 μm and 10 μm, respectively, at a speed of 3 μm/s. We analyzed the force-distance curves using the Hertz model [47], with the assumption that the PA hydrogel is linearly elastic.

### Stem cell culture and cardiomyocyte differentiation

For this work, we used human induced pluripotent stem cell-derived cardiomyocytes (hiPSC-CMs). The hiPSCs were GFP-tagged alpha-actinin-2 (cell line 75) developed at the Allen Institute for Cell Science (allencell.org/cell-catalog) and available through Coriell (AICS-0075-085) [48, 49]. We cultured the hiPSCs on tissue culture plastic coated in Matrigel (Corning, 356252) using feeder-free culture conditions in standard conditions of 5% carbon dioxide at 37°C. The hiPSCs were cultured in Essential 8 Medium (Gibco, ThermoFisher, A1517001) which was changed daily. We passaged the cells with EDTA when confluency reached 75%. We differentiated the hiPSCs into cardiomyocytes (hiPSC-CMs) using a previously published protocol [50] and maintained the hiPSC-CMs until seeding in RPMI 1640 Medium (ThermoFisher, 11875119) with B-27 Supplement (B27; ThermoFisher, 17504044). We seeded the hiPSC-CMs on devices at a density of ∼100,000 cells/cm^2^ between day 24-30 and imaged 3-4 days after seeding (day 27-34).

### Microscopy

We performed all microscopy with a Zeiss Axio Observer 7 inverted microscope with a high speed camera (Photometrics Prime 95b) and a water immersion 40X objective (Plan Apochromat, 1.2 NA). The microscope was equipped with an incubation chamber (PeCon) that maintained a temperature of 37°C and 5% CO_2_ during live-cell imaging.

Before live-cell imaging, we changed the media from B27 to B-27 Supplement in RPMI 1640 Medium, no phenol red (ThermoFisher, 11835030) with 10mM HEPES and 1% Anti-Anti. For each cell, we took a still image in brightfield, 405 (fluorescent microbeads), 488 (alpha-actinin), and 546 (Protein A). We used the still images in the 546 channel to determine whether the cell was on a laminin-only or dual-protein pattern. After the still images, we recorded ∼10 second long videos of sarcomeres (488 channel) and fluorescent microbeads (405 channel). The frame rate was ∼40 frames per second for the sarcomere videos and ∼80 frames per second for the microbeads videos. For the data presented in this work, unless otherwise noted, we included only cells that overlapped at least one N-cadherin end-cap in the dual-protein datasets.

### Sarcomere contractility quantification

We acquired sarcomere shortening videos using alpha-actinin-tagged hiPSC-CMs (as described in *Methods)*. After collecting a video, we cropped it in FIJI (ImageJ) [51] and applied the “Subtract Background” tool with a rolling ball radius of 5 pixels. We then adjusted the brightness and contrast in FIJI using the “auto” option and saved the video as an AVI with a 40 frames per second frame rate.

To quantify the average alignment, sarcomere length, sarcomere shortening, and radial contraction, we ran the videos through Sarc-Graph, a previously published, open-source code that segments the images and tracks sarcomere alignment and contraction [31].

### Traction force microscopy

We utilized CONTRAX, a custom, open source workflow that acquires images and videos, then analyses fluorescent bead displacements using an Ncorr tracking module on pairwise frames from videos of beating hiPSC-CMs to generate matrices of spatiotemporal displacement field data [33, 34]. The traction force microscopy (TFM) module in CONTRAX uses the displacements to calculate traction stresses. The traction stresses are then integrated over the area of the cell to determine the total traction force produced by the cell [33].

We first uploaded bead displacement videos (∼800 frames each) and their corresponding brightfield still images into CONTRAX. For each video, we drew the outline of the cell in FIJI using a composite image of the brightfield and 488 still images of the cell. We saved the cell outline as an ROI and loaded it into CONTRAX with the corresponding bead displacement video (Figure 8a). CONTRAX uses the cell outline to isolate the forces produced by the cell from background noise. The size of the area around the cell included in the analysis can be set using the “mask parameters” in CONTRAX.

**Figure 8.**
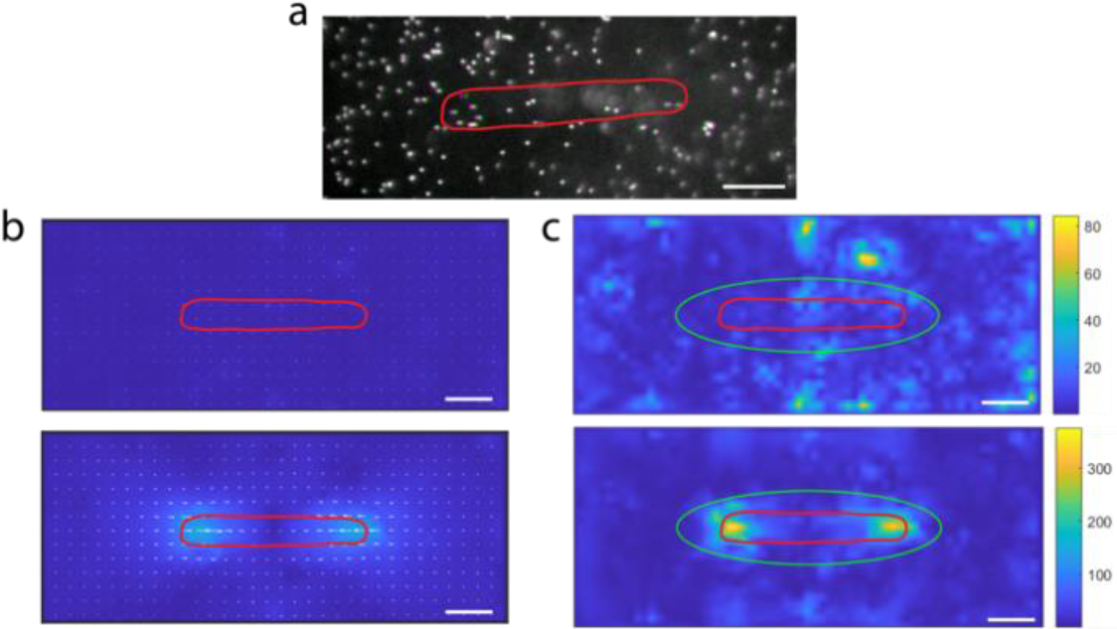
Intermediate steps of traction force microscopy. **(a)** Representative image of fluorescent microbeads when cell is fully relaxed (magenta) and fully contracted (green) state. Representative images of **(b)** displacement maps and **(c)** traction stress maps at relaxed (top) and contracted (bottom) states. Color bars in (c) are in units of Pascals. In all images, red elliptical shape is cell outline. In (c), green ellipse is the analysis region used by CONTRAX. Scale bars are 20 μm. For representative examples of full bead videos, displacement map videos, and traction stress map videos, see supplemental information.

Force production is calculated from bead displacements, so we can expect to see forces slightly outside of the cell outline due to the cell deforming the gel at the cell boundary, causing bead displacements outside the cell outline, as can be seen in Figure 8a.

To capture the forces at the cell boundary, while excluding background noise, we adjusted the “area factor” and “scale factor” in the “mask parameters” section of CONTRAX. After outlining the cell, CONTRAX calculates an ellipse that best fits the outline. The area included in the analysis is equal to the best fit ellipse times the “area factor”, so a larger “area factor” equals a larger analyzed area. The “scale factor” controls the aspect ratio of the analysis region ellipse, with a “scale factor” of greater than 1 increasing the aspect ratio and a “scale factor” of less than 1 decreasing the aspect ratio.

In this work, we chose an analysis region “area factor” of 3 and a “scale factor” of 0.75. The area factor of 3 allows us to capture the forces just outside the cell boundary as described above, while excluding much of the noise around the cell, as can be seen in Figure 8c. The “scale factor” of 0.75 ensures the analysis region stays within the frame, especially with long, thin cells.

After loading the cell outline, we input the material properties for the PA hydrogel the cell was on. For this study, all the cells were cultured on PA hydrogels with a Young’s modulus of ∼6.8 kPa. The Poisson’s ratio for PA hydrogels is generally accepted to be between 0.45 and 0.5 [52–54], and in this work, we used a Poisson’s ratio of 0.45. These parameters are used in the force calculation with the assumption that the gel is linearly elastic and homogenous, which is a generally accepted approximation when the strain on the hydrogel is very small (less than 1), as it is in TFM [53]. Additionally, we assume that the gel is thick enough and wide enough to prevent the cells from experiencing any boundary effects of the gel and that all cell-produced forces normal to the hydrogel surface are negligible. These assumptions are approximately true given that the hydrogel is ∼10 times as thick as the cell height [55] and are generally accepted assumptions for traction force microscopy [53, 55, 56]. After the video initialization, we set the displacement parameters for the Ncorr displacement tracking module. We used a subset radius of 30 px, spacing coefficient of 10 px, cutoff norm of 1e-6, and cutoff iteration of 20. These parameters are used to optimize the bead tracking in Ncorr, minimizing the amplification of noise and the loss of information due to over-smoothing.

Using the resulting displacement maps at each timepoint (Figure 8b), CONTRAX calculates the regularization parameter, lambda, based on L-curve optimization [57]. The regularization parameter constrains the accepted force vectors, minimizing the error due to noise and the error due over-smoothing. Finally, using this regularization parameter, CONTRAX calculates the traction force heatmaps for each timepoint (Figure 8c) using Fourier Transform Traction Cytometry (FTTC) [36, 58].

### Cell fixation and immunostaining

Following live cell imaging, we fixed the hiPSC-CMs with 4% formaldehyde (ThermoFisher, 28908) for 10 minutes. Then we washed the cells with PBS three times and stored them in PBS at 4°C until immunostaining. Before antibody incubation, we incubated the cells for 5 minutes with a permeabilization solution of 0.1% Triton X-100 (ThermoFisher, A16046.AE) in PBS. We then incubated them with a blocking solution of 0.3% Tween20 (ThermoFisher, 28352) and 2% BSA in PBS for 30 minutes. We diluted a pan-cadherin primary antibody (Sigma, C3678) 1:200 in a solution of 0.1% Tween20 and 1% BSA in PBS (dilution buffer). We incubated the hiPSC-CMs with the pan-cadherin antibody solution for 1 hour at room temperature, then washed three times with dilution buffer. We diluted anti-rabbit AF-647 secondary antibody (ThermoFisher, A-32733) 1:500 in dilution buffer and incubated on the cells for 1 hour at room temperature. We washed the cells three times with PBS and then stored in PBS at 4°C until imaging.

### Statistics

Unless otherwise noted, we determined statistical significance of the data presented using parametric, unpaired, two-tailed T-tests. P-values with significance at P<0.05 are designated with (*), P<0.005 are designated with (**), P<0.0005 are designated with (***), and P<0.0001 are designated with (****). For all parametric T-tests, we verified Normality and Lognormality and transformed data as appropriate. For all T-tests, we also performed F-tests to check differences in variance between the two samples. Unless otherwise noted, all F-tests were not significant. For data with significant differences in variance, we reanalyzed the data with a Welch’s correction to account for varying standard deviations. All statistical analyses were performed and visualized using Prism (GraphPad Software, Inc.).

## Author Contributions

Kerry V. Lane: conceptualization, data curation, formal analysis, investigation, methodology, project administration, validation, visualization, writing – original draft, and writing – review & editing. Liam P. Dow: conceptualization, investigation, methodology, visualization, and writing – review & editing. Erica A. Castillo: conceptualization, methodology, and writing – review & editing. Rémi Boros: formal analysis and resources. Sam Feinstein: resources and writing – review & editing. Gaspard Pardon: conceptualization, software, and writing – review & editing. Beth L. Pruitt: conceptualization, funding acquisition, resources, supervision, and writing – review & editing.

## Acknowledgements

This work was supported by National Institutes of Health grant 3RM1GM131981-02S1 (B.L.P.). K.V.L. was supported by NSF GRFP and UCSB fellowship funding. E.A.C. was supported by NSF GFRP and Ford Foundation Predoctoral Fellowship. R.B. was supported by NSF-DMR-2004617.

K.V.L. acknowledges useful conversations with Janae Gayle about the Sarc-Graph code. K.V.L. acknowledges support in hiPSC-CM culture from Trevor Pyle and Danny Gillissen. The authors acknowledge the use of the Nanostructures Cleanroom Facility within the California NanoSystems Institute, supported by the University of California, Santa Barbara and the University of California, Office of the President.

## References

1. Yang, X., L. Pabon, and C.E. Murry, Engineering adolescence: maturation of human pluripotent stem cell-derived cardiomyocytes. Circ Res, 2014. 114(3): p. 511–23.

2. Chen, V.C., et al., Development of a scalable suspension culture for cardiac differentiation from human pluripotent stem cells. Stem Cell Res, 2015. 15(2): p. 365–75.

3. Sayed, N., C. Liu, and J.C. Wu, Translation of Human-Induced Pluripotent Stem Cells: From Clinical Trial in a Dish to Precision Medicine. J Am Coll Cardiol, 2016. 67(18): p. 2161–2176.

4. Schroer, A., et al., Engineering hiPSC cardiomyocyte in vitro model systems for functional and structural assessment. Progress in Biophysics and Molecular Biology, 2019. 144: p. 3–15.

5. Zhang, J., et al., Functional cardiomyocytes derived from human induced pluripotent stem cells. Circulation Research, 2009. 104(4).

6. Ahmed, R.E., et al., A Brief Review of Current Maturation Methods for Human Induced Pluripotent Stem Cells-Derived Cardiomyocytes. Front Cell Dev Biol, 2020. 8: p. 178.

7. Eschenhagen, T., et al., Three-dimensional reconstitution of embryonic cardiomyocytes in a collagen matrix: a new heart muscle model system. The FASEB Journal, 1997. 11(8): p. 683–694.

8. Tulloch, N.L., et al., Growth of Engineered Human Myocardium With Mechanical Loading and Vascular Coculture. Circulation Research, 2011. 109(1): p. 47–59.

9. Mannhardt, I., et al., Human Engineered Heart Tissue: Analysis of Contractile Force. Stem Cell Reports, 2016. 7(1): p. 29–42.

10. Ronaldson-Bouchard, K., et al., Advanced maturation of human cardiac tissue grown from pluripotent stem cells. Nature, 2018. 556(7700): p. 239–243.

11. Sager, P.T., et al., Rechanneling the cardiac proarrhythmia safety paradigm: a meeting report from the Cardiac Safety Research Consortium. Am Heart J, 2014. 167(3): p. 292–300.

12. Crumb, W.J., Jr., et al., An evaluation of 30 clinical drugs against the comprehensive in vitro proarrhythmia assay (CiPA) proposed ion channel panel. J Pharmacol Toxicol Methods, 2016. 81: p. 251–62.

13. Fermini, B., et al., A New Perspective in the Field of Cardiac Safety Testing through the Comprehensive In Vitro Proarrhythmia Assay Paradigm. J Biomol Screen, 2016. 21(1): p. 1–11.

14. Blair, C.A. and B.L. Pruitt, Mechanobiology Assays with Applications in Cardiomyocyte Biology and Cardiotoxicity. Adv Healthc Mater, 2020. 9(8): p. e1901656.

15. Wang, G., et al., Modeling the mitochondrial cardiomyopathy of Barth syndrome with induced pluripotent stem cell and heart-on-chip technologies. Nature Medicine, 2014. 20(6): p. 616–623.

16. Ribeiro, A.J., et al., Contractility of single cardiomyocytes differentiated from pluripotent stem cells depends on physiological shape and substrate stiffness. Proc Natl Acad Sci U S A, 2015. 112(41): p. 12705–10.

17. Bray, M.-A., S.P. Sheehy, and K.K. Parker, Sarcomere alignment is regulated by myocyte shape. Cell Motility and the Cytoskeleton, 2008. 65(8): p. 641–651.

18. Thery, M., et al., Anisotropy of cell adhesive microenvironment governs cell internal organization and orientation of polarity. Proc Natl Acad Sci U S A, 2006. 103(52): p. 19771–6.

19. Tseng, Q., et al., Spatial organization of the extracellular matrix regulates cell-cell junction positioning. Proceedings of the National Academy of Sciences, 2012. 109(5): p. 1506–1511.

20. Rothenberg, K.E., et al., Controlling Cell Geometry Affects the Spatial Distribution of Load Across Vinculin. Cellular and Molecular Bioengineering, 2015. 8(3): p. 364–382.

21. Chopra, A., et al., Cardiac myocyte remodeling mediated by N-cadherin-dependent mechanosensing. American Journal of Physiology - Heart and Circulatory Physiology, 2011. 300(4): p. 1252–1266.

22. Chopra, A., et al., α-Catenin Localization and Sarcomere Self-Organization on N-Cadherin Adhesive Patterns Are Myocyte Contractility Driven. PLoS ONE, 2012. 7(10).

23. Milani-Nejad, N. and P.M.L. Janssen, Small and large animal models in cardiac contraction research: Advantages and disadvantages. Pharmacology & Therapeutics, 2014. 141(3): p. 235–249.

24. Hasenfuss, G., Animal models of human cardiovascular disease, heart failure and hypertrophy. Cardiovascular Research, 1998. 39(1): p. 60–76.

25. Goncharova, E.J., Z. Kam, and B. Geiger, The involvement of adherens junction components in myofibrillogenesis in cultured cardiac myocytes. Development, 1992. 114(1): p. 173–183.

26. Simpson, D.G., et al., Contractile activity and cell-cell contact regulate myofibrillar organization in cultured cardiac myocytes. Journal of Cell Biology, 1993. 123(2): p. 323–336.

27. Wu, J.C., et al., Role of N-cadherin- and integrin-based costameres in the development of rat cardiomyocytes. J Cell Biochem, 2002. 84(4): p. 717–24.

28. Loh, C.Y., et al., The E-Cadherin and N-Cadherin Switch in Epithelial-to-Mesenchymal Transition: Signaling, Therapeutic Implications, and Challenges. Cells, 2019. 8(10): p. 1118–1118.

29. Sarker, B., C. Walter, and A. Pathak, Direct Micropatterning of Extracellular Matrix Proteins on Functionalized Polyacrylamide Hydrogels Shows Geometric Regulation of Cell-Cell Junctions. ACS Biomaterials Science and Engineering, 2018. 4(7): p. 2340–2349.

30. Nag, K., et al., Cadherin-Fc Chimeric Protein-Based Biomaterials: Advancing Stem Cell Technology and Regenerative Medicine Towards Application, C. Atwood and S.V. Meethal, Editors. 2014. p. 137–164.

31. Zhao, B., et al., Sarc-Graph: Automated segmentation, tracking, and analysis of sarcomeres in hiPSC-derived cardiomyocytes. PLoS Comput Biol, 2021. 17(10): p. e1009443.

32. McCain, M.L., et al., Cooperative coupling of cell-matrix and cell-cell adhesions in cardiac muscle. Proceedings of the National Academy of Sciences of the United States of America, 2012. 109(25): p. 9881–9886.

33. Pardon, G., et al., Insights into single hiPSC-derived cardiomyocyte phenotypes and maturation using ConTraX, an efficient pipeline for tracking contractile dynamics. 2021, Cold Spring Harbor Laboratory.

34. Pardon, G., ContraX. 2021: GitHub repository.

35. Ribeiro, A.J.S., et al., Multi-imaging method to assay the contractile mechanical output of micropatterned human iPSC-derived cardiac myocytes. Circulation Research, 2017. 120(10): p. 1572–1583.

36. Butler, J.P., et al., Traction fields, moments, and strain energy that cells exert on their surroundings. Am J Physiol Cell Physiol, 2002. 282(3): p. C595–605.

37. Pasqualini, F.S., et al., Traction force microscopy of engineered cardiac tissues. PLoS One, 2018. 13(3): p. e0194706.

38. Wheelwright, M., et al., Investigation of human iPSC-derived cardiac myocyte functional maturation by single cell traction force microscopy. PLoS One, 2018. 13(4): p. e0194909.

39. Davis, J., et al., A Tension-Based Model Distinguishes Hypertrophic versus Dilated Cardiomyopathy. Cell, 2016. 165(5): p. 1147–1159.

40. Gavard, J., et al., Lamellipodium extension and cadherin adhesion: two cell responses to cadherin activation relying on distinct signalling pathways. J Cell Sci, 2004. 117(Pt 2): p. 257–70.

41. Collins, C., et al., Changes in E-cadherin rigidity sensing regulate cell adhesion. Proceedings of the National Academy of Sciences of the United States of America, 2017. 114(29): p. E5835–E5844.

42. Goding, J.W., Use of Staphylococcal Protein-a as an Immunological Reagent. Journal of Immunological Methods, 1978. 20(Apr): p. 241–253.

43. Drees, F., A. Reilein, and W.J. Nelson, Cell-Adhesion Assays, in Cell Migration. Methods in Molecular Biology, J.L. Guan, Editor. 2005, Humana Press.

44. Edelstein, A.D., et al., Advanced methods of microscope control using muManager software. J Biol Methods, 2014. 1(2).

45. Denisin, A.K. and B.L. Pruitt, Tuning the Range of Polyacrylamide Gel Stiffness for Mechanobiology Applications. ACS Applied Materials and Interfaces, 2016. 8(34): p. 21893–21902.

46. Van Vliet, K.J., Instrumentation and Experimentation, in Handbook of Nanoindentation with Biological Applications, M.L. Oyen, Editor. 2019, Jenny Stanford Publishing. p. 39−76.

47. Lin, D.C. and F. Horkay, Nanomechanics of polymer gels and biological tissues: A critical review of analytical approaches in the Hertzian regime and beyond. Soft Matter, 2008. 4(4): p. 669–682.

48. Roberts, B., et al., Systematic gene tagging using CRISPR/Cas9 in human stem cells to illuminate cell organization. Molecular Biology of the Cell, 2017. 28(21): p. 2854–2874.

49. Roberts, B., et al., Fluorescent Gene Tagging of Transcriptionally Silent Genes in hiPSCs. Stem Cell Reports, 2019. 12(5): p. 1145–1158.

50. Sharma, A., et al., Derivation of highly purified cardiomyocytes from human induced pluripotent stem cells using small molecule-modulated differentiation and subsequent glucose starvation. Journal of Visualized Experiments, 2015. 2015(97): p. 52628–52628.

51. Schindelin, J., et al., Fiji: an open-source platform for biological-image analysis. Nat Methods, 2012. 9(7): p. 676–82.

52. Kandow, C.E., et al., Polyacrylamide Hydrogels for Cell Mechanics: Steps Toward Optimization and Alternative Uses. Methods in Cell Biology, 2007. 83(07): p. 29–46.

53. Schwarz, U.S. and J.R. Soine, Traction force microscopy on soft elastic substrates: A guide to recent computational advances. Biochim Biophys Acta, 2015. 1853(11 Pt B): p. 3095–104.

54. Dembo, M. and Y.L. Wang, Stresses at the cell-to-substrate interface during locomotion of fibroblasts. Biophysical Journal, 1999. 76(4): p. 2307–2316.

55. Kraning-Rush, C.M., et al., Quantifying Traction Stresses in Adherent Cells. 2012, Academic Press Inc. p. 139–178.

56. Ribeiro, A.J., et al., For whom the cells pull: Hydrogel and micropost devices for measuring traction forces. Methods, 2016. 94: p. 51–64.

57. Hansen, P.C. and D.P. O’Leary, The Use of the L-Curve in the Regularization of Discrete Ill-Posed Problems. SIAM Journal on Scientific Computing, 1993. 14(6): p. 1487–1503.

58. Kulkarni, A.H., et al., Traction cytometry: regularization in the Fourier approach and comparisons with finite element method. Soft Matter, 2018. 14(23): p. 4687–4695.

