## Supplemental Figures 1-4 for "Spatial Patterning of Laminin and N-Cadherin for Human Induced Pluripotent Stem Cell-Derived Cardiomyocytes (hiPSC-CMs)"

### **Supplemental Information**

To verify the N-cadherin patterning, we made single-protein patterns with N-cadherin on glass coverslips via lift-off protein patterning [1]. We stained the devices with a pan-cadherin antibody (Sigma, C3678) and AF-488 secondary (ThermoFisher, A-11034; Figure S1a). We tested the specificity of the pan-cadherin antibody with a number of controls: 1) no protein with pan-cadherin primary and AF-488 secondary antibodies (Figure S1b), 2) no protein with AF-488 secondary antibody (Figure S1c), 3) laminin with pan-cadherin primary and AF-488 secondary antibodies (Figure S1d), and 4) Protein A with pan-cadherin primary and AF-488 secondary antibodies (Figure S1e). For each control, we performed lift-off patterning on a glass coverslip before incubating with protein or antibody, which is why we still see the patterns in the control experiments (Figure S1).

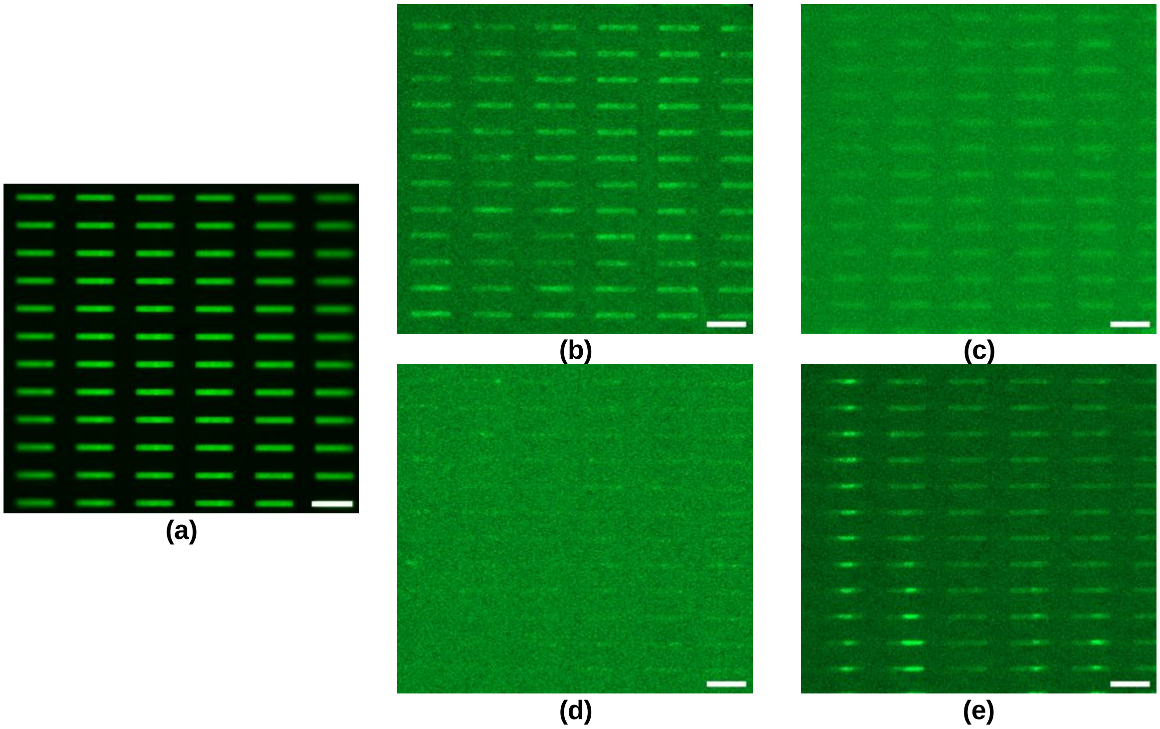


**Figure S1. Control experiments demonstrated specificity of pan-cadherin antibody.** Fluorescence images of **(a)** N-cadherin protein pattern on glass, and immunostaining controls on glass: **(b)** no protein devices stained with pan-cadherin and AF488 antibodies, **(c)** no protein devices stained with AF488 antibody, **(d)** devices patterned with laminin stained with pan-cadherin and AF488 antibodies, and **(e)** devices patterned with Protein A stained with pan-cadherin and AF488 antibodies. Scale bars are 100 μm.

#### hiPSC-CMs Did Not Attach to N-cadherin-Patterned PA Hydrogels without oHEA Functionalization

In preliminary experiments, we seeded hiPSC-CMs on single-protein N-cadherin patterns on both glass and unfunctionalized PA hydrogels (Figure S2). We found that hiPSC-CMs would attach to N-cadherin patterns on glass but not to N-cadherin patterns on PA hydrogels without oHEA functionalization (Figure S2c-d). To rule out a problem with the PA hydrogel, we also made single-protein patterns with Matrigel (Corning, 356252), a basement membrane protein cocktail that hiPSC-CMs are known to attach to [2]. We saw that hiPSC-CMs readily attached to Matrigel patterns on both glass and unfunctionalized PA hydrogels (Figure S2a-b).

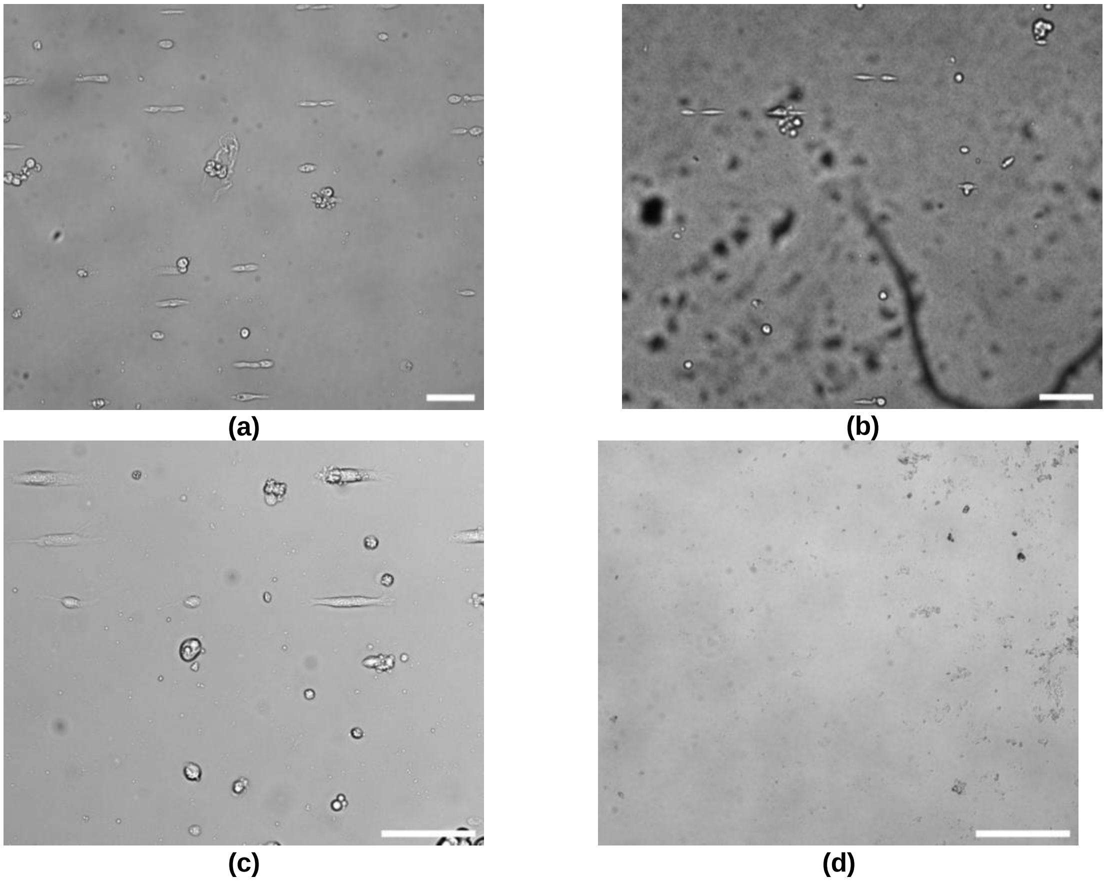


**Figure S2.** hiPSC-CMs attach to Matrigel on glass and PA hydrogel, and N-cadherin on glass, but not N-cadherin on PA hydrogel. Brightfield images of hiPSC-CMs seeded on **(a)** Matrigel patterned on glass, **(b)** Matrigel patterned on PA hydrogel, **(c)** N-cadherin patterned on glass, and **(d)** N-cadherin patterned on PA hydrogel. Scale bars are 100 μm.

Because the hiPSC-CMs readily attached to N-cadherin patterns on glass but not PA hydrogels, we hypothesized that N-cadherin was not stably anchored on the unfunctionalized PA hydrogel surface. We stained previously-seeded N-cadherin-patterned PA hydrogels with pan-cadherin primary and AF-488 secondary antibodies

(Figure S3). We found that all N-cadherin patterns had been removed from the hydrogel surface, suggesting that N-cadherin is detached in the process of hiPSC-CM attachment (Figure S3).

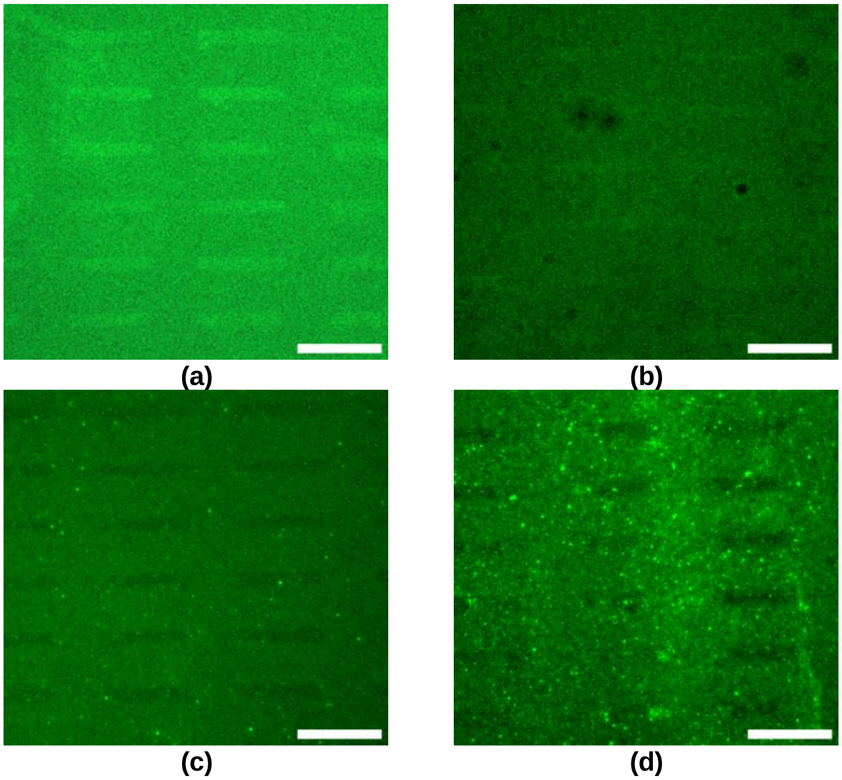


**Figure S3. N-cadherin is removed from PA hydrogel devices after hiPSC-CMs are seeded and fail to attach.** GFP images of N-cadherin-patterned **(a)** glass device 1, **(b)** glass device 2, **(c)** PA hydrogel device 1, and **(d)** PA hydrogel device 2 stained four days after they were made and three days after seeding them with hiPSC-CMs. Patterns were visualized with pan-cadherin primary antibody and AlexaFluor 488 secondary antibody. Scale bars are 100 μm.

To demonstrate that the degradation or removal of N-cadherin is not due to the time elapsed between incubation and staining, we did a time study with N-cadherin-patterned PA hydrogels. We made three N-cadherin-patterned PA hydrogels to stain for N-cadherin patterns at different time points after being cultured in media. The first PA hydrogel was stained and imaged the day of seeding (Figure S4a), the second hydrogel two days after seeding (Figure S4b), and the third hydrogel three days after seeding (Figure S4c). With the unseeded devices, there was slight degradation of the N-cadherin pattern with time, but overall the pattern was preserved, indicating that media and/or time would not result in the N-cadherin removal seen in Figure S3.

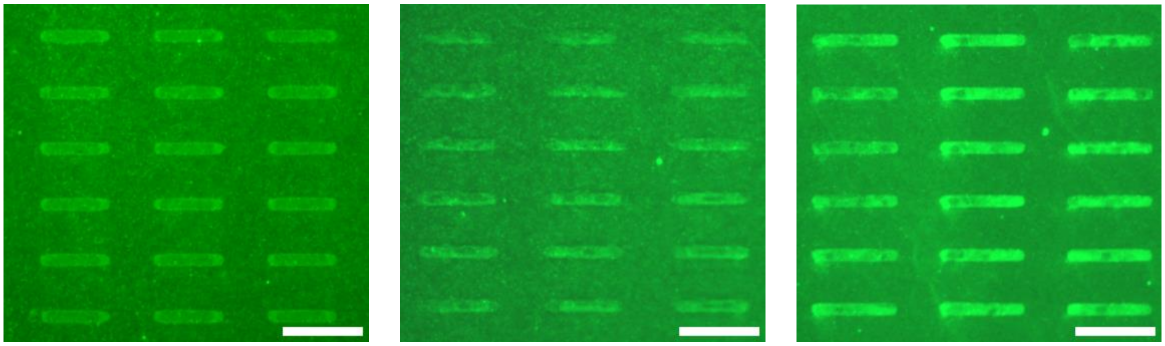

**(a)** **(b)** **(c)**

**Figure S4.** N-cadherin patterns do not degrade over time when cultured in hiPSC-CM media. GFP images of N-cadherin patterned on PA hydrogel devices immunostained **(a)** the day after the devices were made, **(b)** three days after the devices were made, and **(c)** four days after the devices were made. Patterns were visualized with pan-cadherin primary antibody and AF488 secondary antibody. Scale bars are 100 μm.


### References

1. Moeller, J., et al., *Controlling cell shape on hydrogels using liftoff protein patterning.* PLoS ONE, 2018. **13**(1): p. 1-17.
2. Hughes, C.S., L.M. Postovit, and G.A. Lajoie, *Matrigel: A complex protein mixture required for optimal growth of cell culture.* PROTEOMICS, 2010. **10**(9): p. 1886-1890.
